# Cocaine- and amphetamine-regulated transcript in perciforms II. Responsiveness to energetic shortage

**DOI:** 10.64898/2025.12.04.692351

**Authors:** Nikko Alvin R. Cabillon, Liat Koch, Noam Mizrahi, Adi Segev-Hadar, Iris Meiri-Ashkenazi, Gertrude Aluppo, Jakob Biran, Amir Bitan

## Abstract

Cocaine and amphetamine-regulated transcript (Cart) is a neuropeptide with prominent roles in appetite regulation and maintenance of energy homeostasis. Although Cart has been widely studied in vertebrates, its multigenic nature in fish elicits questions regarding the various functions affected by specific cart peptides. This is further emphasized when considering the high variation of aquatic ecosystems fish occupy. Nile tilapia (*Oreochromis niloticus*) and gilthead seabream (*Sparus aurata*) are important aquaculture species with different natural habitats, food preferences and feeding behaviors. Herein, we utilize these two species to evaluate whether food-related cart functions are species-specific. To this end, we studied how short-term (SD) or long-term (LD) food deprivation affects the central expression of the multiple *cart* genes of both species. Quantitative PCR analysis of *cart* expression demonstrated that SD resulted in decreased midbrain expression of three tilapia *carts* and three seabream *carts* while LD decreased the midbrain expression of two tilapia carts and two seabream carts. In addition, SD increased the expression of tilapia *oncart1c* in the anterior brain and *oncart1c* and *oncart1b* in the posterior brain while reducing the expression of seabream *sacart1c*. Our analyses showed that there are *cart* genes in each species (*cart1a* and *cart1b* for tilapia, and *cart1b* for seabream) that responded to both SD and LD by reduced expression in the midbrain. In addition, both conditions reduced the expression of seabream *sacart1c* in the posterior brain. Taken together, our current findings suggest that the major appetite regulation in each species is mediated by species-specific *carts*.

## 1 Introduction

Cocaine and amphetamine-regulated transcript (Cart) is a highly conserved neuropeptide among vertebrates, with an abundant mRNA expression in the hypothalamic region (Elias et al., 2001; Farzi et al., 2018; Kristensen et al., 1998; Vrang, 2006). Cart has been implicated in diverse neuronal functions including stress, anxiety, neuroprotection, addiction, and learning. Nevertheless, the most prominent functions of Cart are the central regulation of feeding and energy homeostasis (Ahmadian-Moghadam et al., 2018; Bharne et al., 2013; Chaki et al., 2003; Conde-Sieira et al., 2018; Lau and Herzog, 2014; Vicentic and Jones, 2007). Several studies in various vertebrate models have shown that food deprivation decreased *cart* mRNA expression in the brain while refeeding resulted in subsequent increase in expression (Lartigue et al., 2007; Zhang et al., 2018; Zhou et al., 2019). Furthermore, cart was shown to actively suppress feeding behavior in more than 20 fish species (Volkoff, 2016). Mammalian genomes contain a single *cart* gene, yet multiple *cart* genes were identified in avian and piscine genomes (Cai et al., 2015; Gutierrez-Ibanez et al., 2016; Soengas et al., 2018). Appetite regulation by cart was suggested to be species-specific and dependent on feeding habits (Butt et al., 2019; Polakof et al., 2012). The expression of cart genes, in response to food deprivation, was studied in several species. In chicken (*Gallus gallus*), food deprivation for 48h resulted in suppressed *cart1* expression, and did not affect *cart2* expression (Cai et al., 2015). In medaka (*Oryzias latipes*), only one out of six *cart* genes decreased in expression after 17 days of food deprivation (Murashita and Kurokawa, 2011). A few studies in fish did not find altered *cart* expression in response to fasting (MacDonald and Volkoff, 2009a, 2009b; Volkoff et al., 2016). The genomes of these species have not been fully sequenced, and it is possible that some of their *cart* genes including the appetite-related *cart* have yet been identified.

The regulation of feed intake and metabolic rate by various neurotransmitters, neuropeptides and hormonal systems was extensively studied in fish (Soengas et al., 2018; Volkoff, 2016). This regulation involves two main components: (1) a short-term ‘meal to meal’ component which is comprised of central and peripheral signals of the brain-gastrointestinal axis, and (2) a long-term component of days to months that is also influenced by humoral feedback from energetic stores (Soengas et al., 2018). In mammals, Cart neurons from the lateral hypothalamus and the hypothalamic arcuate nucleus govern the overall regulation of energy balance (Farzi et al., 2018; Lau et al., 2018). These regions are proximate to the semi-permeable blood-brain barrier thus allowing exposure of the hypothalamic Cart neurons to circulating humoral and metabolic mediators (Barsh and Schwartz, 2002; Berthoud, 2002; Faouzi et al., 2007; Lau and Herzog, 2014). In fish, the nucleus lateralis tuberis (NLT) is homologous to the mammalian arcuate nucleus and abundant *cart* expression has been localized in this brain region in zebrafish (*Danio rerio*) (Akash et al., 2014). This study has further demonstrated that 48 h fasting resulted in the decrease in expression of *cart* in the NLT as well as in the nucleus recessus lateralis and the entopeduncular nucleus, which were previously shown to be involved in energy homeostasis (Akash et al., 2014). Furthermore, there is evidence showing that food deprivation longer than 48 h caused a metabolic disturbance in fish and thus affected brain and liver energetic processing related to the long-term component of feeding (Figueroa et al., 2000). The temporal aspects of feeding regulation may also be bidirectional. For example, when tested in Siberian sturgeon (*Acipenser baerii*), *cart* expression showed an opposite trend in expression at 24 h compared to extended fasting of up to 15 days (Zhang et al., 2018). Nile tilapia and gilthead seabream have fully sequenced genomes that allow deep examination of their multigenic cart systems and their possible involvement in feeding regulation. Nile tilapia is a euryhaline fish that is mainly herbivorous and feed on algae (Levina et al., 2021; Njiru et al., 2004), while gilthead seabream is mostly carnivorous marine species that mainly feed on crustacean, mollusks and small fishes (Mhalhel et al., 2023). Herein, we use these two species to study if and how natural differences in feeding behavior and diet composition affect their regulatory cart system.

We recently identified seven *cart* genes in tilapia and six *cart* genes in seabream and found that their main site of expression is in the midbrain where the hypothalamus is located (Cabillon et al. 2025). Which of the newly identified *cart* genes may have a role in appetite regulation and its responsiveness to the fish metabolic state remained to be tested. Aiming to identify appetite-related *cart*s, tilapia and seabream were subjected to either short-term (SD) or long-term (LD) food deprivation and *cart* responsiveness was analyzed by quantitative-PCR (qPCR). We found that only specific *cart* genes responded to either or both challenges, and that while the expression of some carts is suppressed by SD or LD, expression of other carts increased in response to these metabolic challenges. Our current findings support the existence of main anorexigenic *cart* genes in each species and a functional partitioning in the piscine cart system.

## 2 Methods

### 2.1 Animal Husbandry

The experiments were approved by the Agricultural Research Organization (A.R.O.) Committee for Ethics in Using Experimental Animals (approval number: 877/20). Tilapia samples for LD analyses were taken from the study of (Segev-Hadar et al., 2020) Briefly, the setup was divided into two treatments: one fed twice daily and the other was not fed for 21 days. For SD analyses, adult tilapia (77.93 ± 2.6) were acclimated to experimental tanks 2 weeks prior to the experiment. During acclimation, all fish were fed twice daily *ad libitum* on commercial tilapia feeds containing 33% protein (Zemach feed mill™, Israel). Temperature was maintained at 26°C and physico-chemical parameters were monitored. Feed was last administered to the starved group 24h before harvest. The fed group was maintained under regular feeding regime. For seabream, fish (92.37± 6.06 g) were acclimated for 9 days in sea water (with salinity of about 40‰) open flow through tanks (270L), under natural ambient temperature in the range of 22-24°C and divided for 3 treatments (3 tanks per treatment) as follows: fed fish that were fed twice a day (Raanan food, Israel) with food containing 46% protein *ad libitum*, or fish that were food deprived for 48 h or 7 days prior to sampling.

### 2.2 Tissue collection and RNA extraction

Fish were subsequently euthanized and brains were harvested. Piscine hypothalamic nuclei are localized in the diencephalic compartment (Faouzi et al., 2007). To gain better resolution of the hypothalamic-related CART expression, the midbrain section containing the diencephalon and optic tectum was microdissected from other brain compartments, placed in a tube, snap-frozen in liquid nitrogen and stored at -80°C until further analysis.

Total RNA was extracted using a trizol/chloroform extraction with Tri-reagent (BioLab, Jerusalem, Israel) for seabream and using Trizol reagent (Life technologies Corporation, Carlsbad, USA), for tilapia according to the manufacturer’s protocols. Total RNA quantity and purity were measured using a microplate spectrophotometer Epoch™ (BioTek instruments Inc. Winooski, USA) for tilapia or by NanoDropOne (Thermo scientific Paisley, UK) for seabream. Possible genomic DNA contamination was eliminated by treatment with TURBO DNAse-free™ kit Invitrogen (Thermo Fisher Scientific, Vilnius, Lithuania) for tilapia or with PerfeCTa® DNase I RNase-free (Quantabio, Beverly, USA) for seabream according to the manufacturer’s protocols. DNase free total RNA (0.5 µg) was reverse transcribed using High Capacity cDNA Reverse transcription kit (Thermo Fisher Scientific, Vilnius, Lithuania) for tilapia or qScript cDNA Synthesis Kit (Quantabio, Beverly, USA) for seabream according to the manufacturer’s protocols. The cDNA was stored at -20^0^C until further analysis.

### 2.3 Realtime Quantitative (q)PCR

Gene expression levels were analyzed by qPCR using a StepOnePlus™ Real-Time PCR System (Applied Biosystems, Inc. Foster City, CA, USA). Elongation factor 1 alpha (*ef1α*) and 18S served as reference genes (Malandrakis et al., 2014; Segev-Hadar et al., 2020). For tilapia, each reaction consisted of 5 microliter (µL) SYBR® green dye (Thermo Fisher Scientific, Vilnius, Lithuania), 0.75µL of 3 µM forward and reverse primers, 0.5 µL of Ultra-Pure Water (UPW) and 3µL of cDNA template (all tissues diluted 1:10 in UPW). Analysis was performed in duplicates. Controls without the cDNA were used to test for non-specific amplification. Melt curve analysis was used to confirm amplification of a single product. Amplification was performed under the following conditions; 95°C for 20s, 40 cycles at 95°C for 3s, and 60°C for 30s, followed by one cycle at 95°C for 15s 60°C for 1 min, 95°C for 15s for the generation of the melting curve. For seabream, reactions were prepared in a final volume of 10µL containing 2µL of 50ng cDNA, 0.3µL of each forward and reverse primers (10µM) to total concentration of 3µM, 5µL of the SYBR Green FastMix ROX Kit (Quanta bio) and µL of water. All primers used in this study are detailed in **Table 1**. Reactions were conducted in 96-well plates and samples were analyzed in triplicate. In all cases, for each gene, negative control was included. Expression levels were analyzed by Real time PCR using a StepOnePlus™ System (Applied Biosystems, Inc. Foster City, CA, USA). For both species, relative quantification was determined using the 2^−ΔΔCT^ method (Rao et al., 2013), with fed fish group as controls.

**Table 1.**
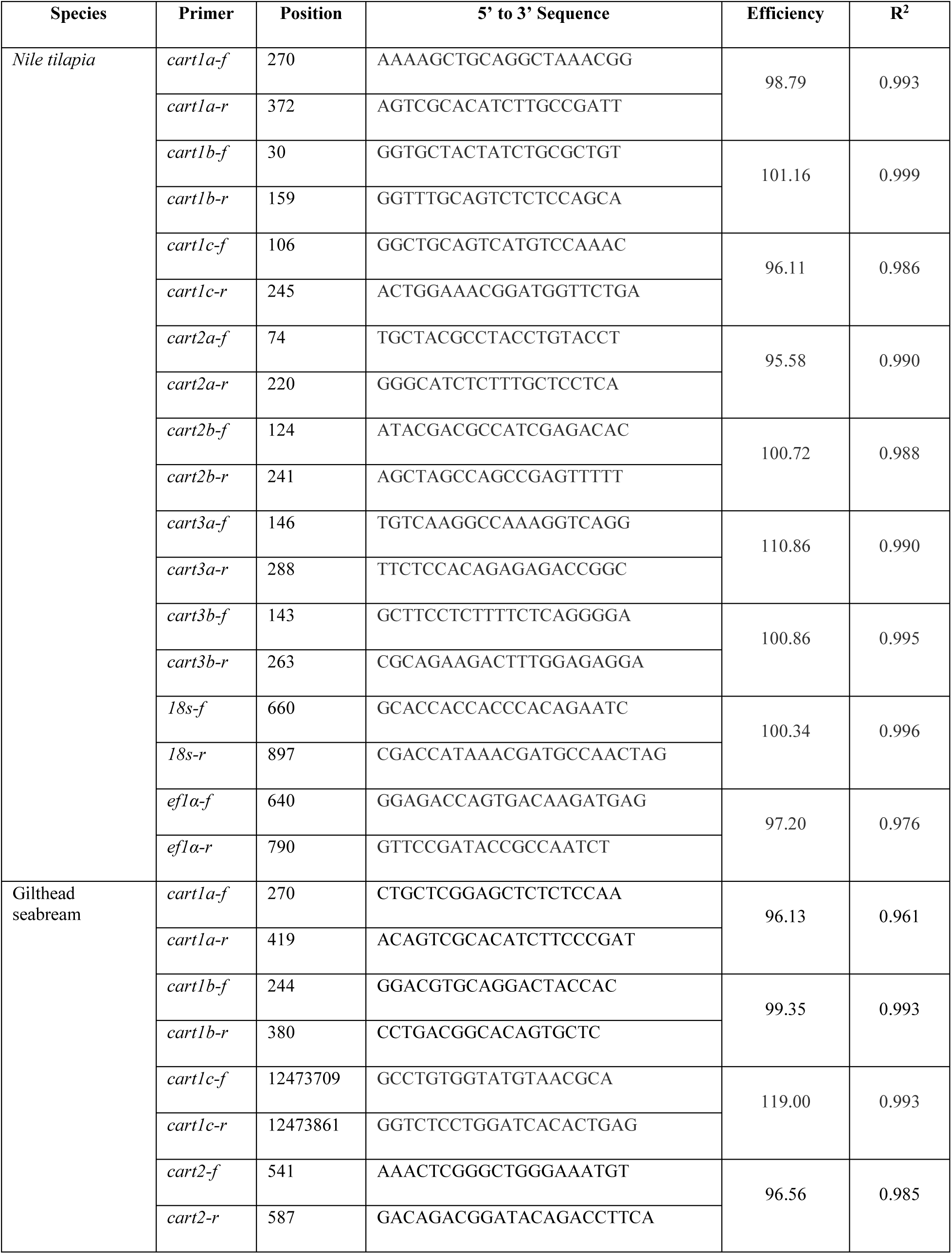
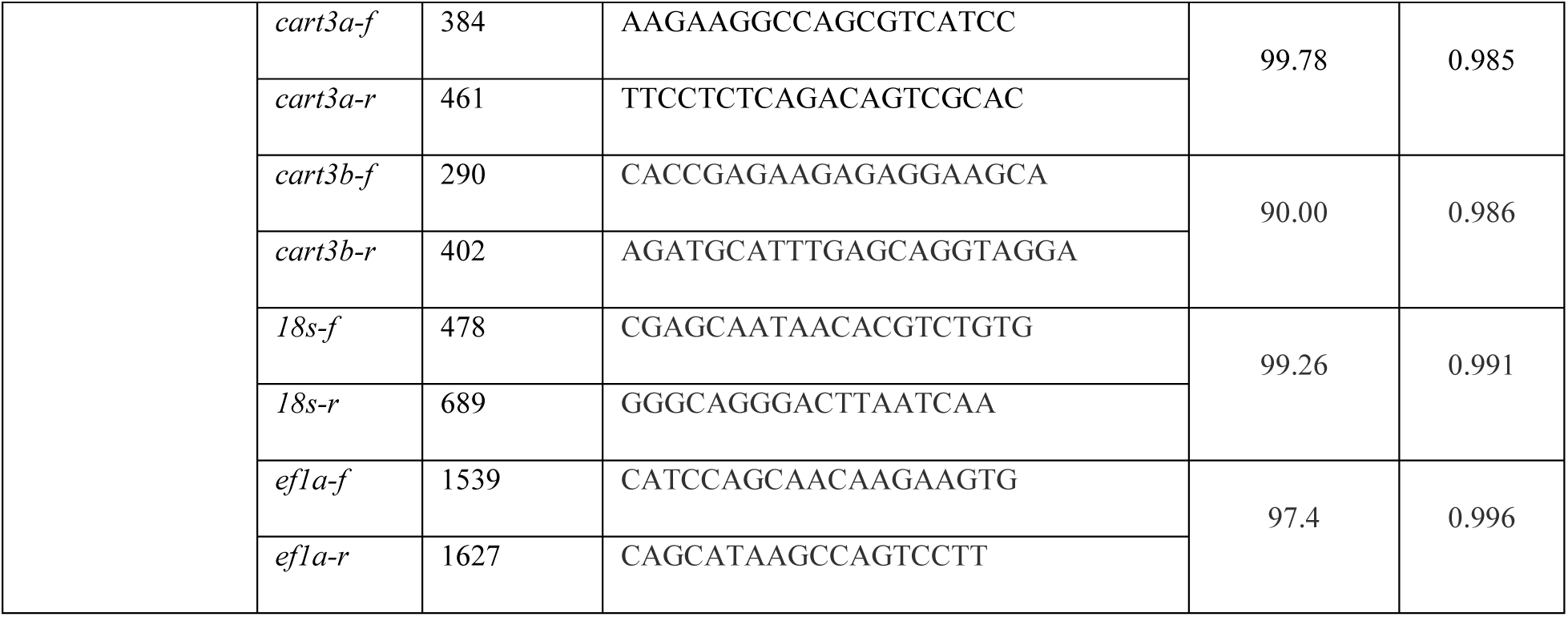
Primer sets for quantitative PCR.

### 2.4 Statistical analyses

GraphPad Prism version 8 (GraphPad, San Diego, USA) was used to perform the statistical analyses. Grubb’s test was used to remove outlier. Multiple student’s t-test was employed to identify significant differences in the expression between treated and control groups. The level of significance was set at P < 0.05.

## 3 Results

### 3.1 Differential effects of short-term food deprivation (SD) on carts expression

Cart is mostly known for its role as an anorexigenic neuropeptide. However, in view of its pleiotropic functions, the existence of multiple *cart* genes may result in functional portioning or redundancy as was previously shown in zebrafish (Ahi et al., 2019). Aiming to identify cart genes involved in regulation of feeding and appetite, fish were deprived of food for short time periods and their brains were analyzed for *cart* mRNA expression. Nile tilapia subjected to SD showed a significant decrease in midbrain expression of *oncart1a* and *oncart3a* as well as a decreasing trend in the expression of *oncart1b* (p = 0.08) (**Fig. 1**). *oncart1a* decreased in expression in the anterior brain while *oncart1b* increased in expression in the posterior brain. SD also led to increased expression of *oncart1c* in both anterior and posterior brain regions. Seabream subjected to SD exhibited a significant decrease in midbrain expression of *sacart1b, sacart1c* and *sacart3b* (**Fig. 2**). Significantly decreased *sacart1c* mRNA was also noted in the anterior and posterior brain compartments (**Fig. 2**). These data show that feeding-related cart genes may vary according to the fish species.

**Figure 1.**
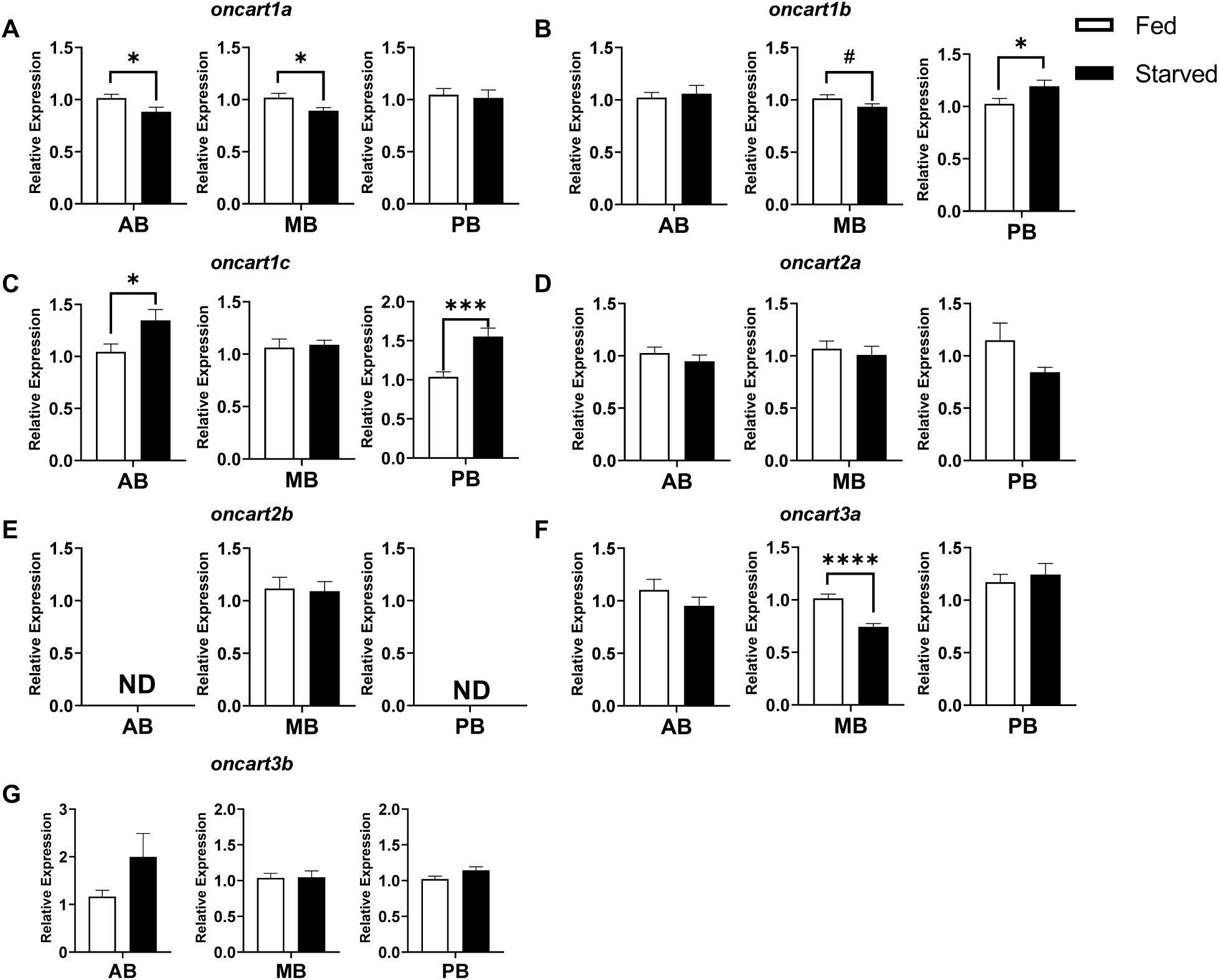
Relative expression of *cart* orthologues in major compartments of Nile tilapia brain following short term food deprivation (SD). Adult tilapia were not fed for 24h. Expression of *oncart1a* (**A**), *oncart1b* (**B**), *oncart1c* (**C**), *oncart2a* (**D**), *oncart2b* (**E**), *oncart3a* (**F**), *oncart3b* (**G**) was analyzed in the anterior region, (AB) midbrain region (MB), and posterior region (PB) of the brain using qPCR (control n = 24, starved n = 24). Significant difference (p < 0.05) is indicated by (*), p <0.01 (**), p <0.001 (***), p <0.0001 (****) and p = 0.087 (#). Multiple t-test with significant value set at p < 0.05.

**Figure 2.**
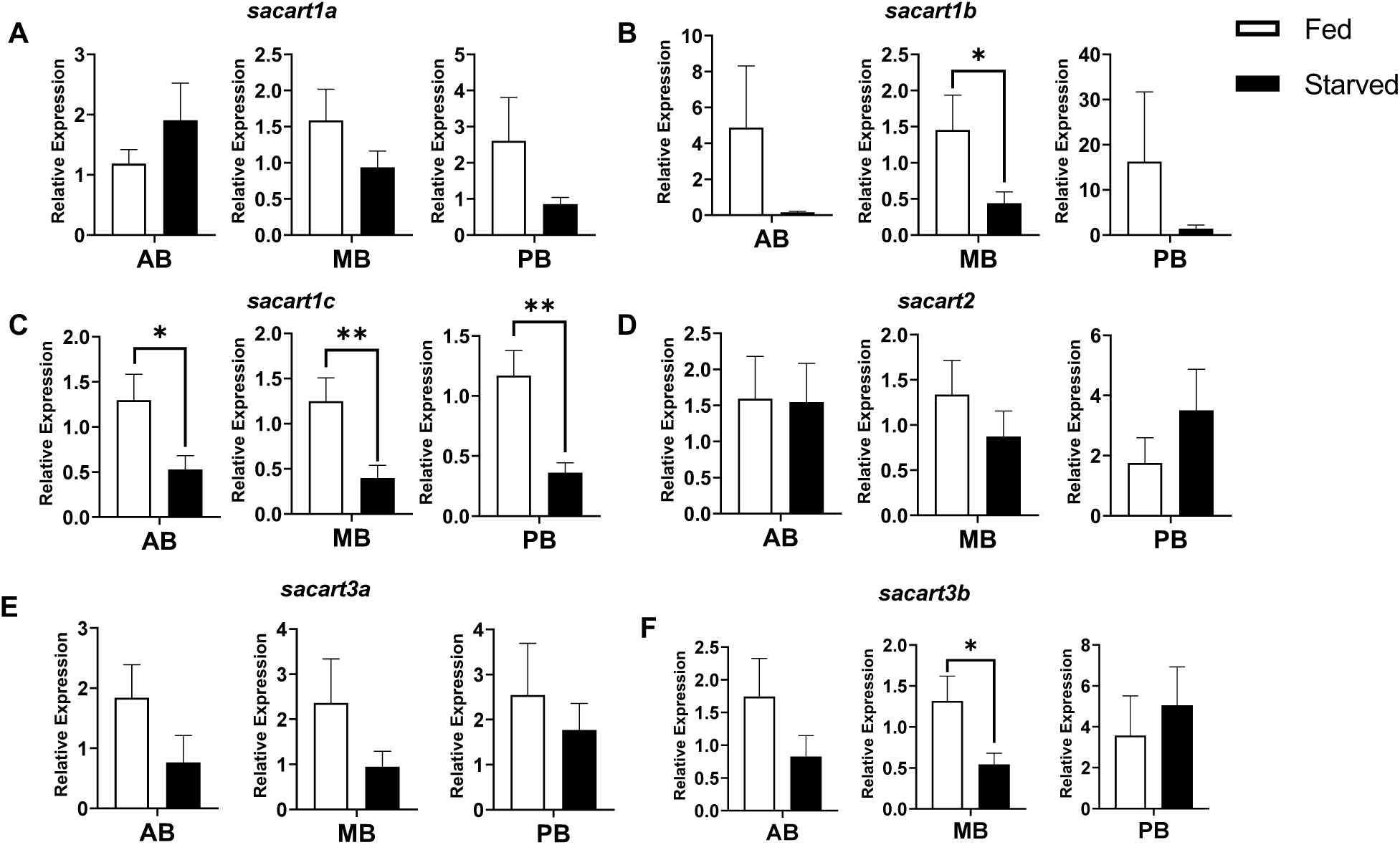
Relative expression of *cart* orthologues in major compartments of sea bream brain following SD. Adult seabream were not fed for 48h. Expression of *sacart1a* (**A**), *sacart1b* (**B**), *sacart1c* (**C**), *sacart2* (**D**), *sacart3a* (**E**), *sacart3b* (**F**) was analyzed in the AB, MB, and PB of the brain using qPCR (control n = 8, starved n = 10). Significant difference (p < 0.05) is indicated by (*), and p <0.01 (**).

### 3.2 Other carts are affected by long-term food deprivation (LD)

Food deprivation for 21 days in Nile tilapia led to a decrease in expression of *oncart1a* and *oncart1b* in the midbrain (**Fig. 3**). In the anterior brain of tilapia, a decrease in expression of *oncart1b* and *oncart3a* was observed. A significant decrease in expression of *oncart2a* was also observed in the posterior brain. Moreover, while *oncart3a* expression significantly decreased in the anterior brain, a significant increase in expression was observed in the posterior brain. In seabream, 7 days of food deprivation resulted in decreased expression of *sacart1b* and a trend (p = 0.070) of decreased expression of *sacart2* in the midbrain (**Fig. 4**). No significant change in expression in the anterior brain was observed in the seabream, but a decreased expression of *sacart1c* was identified in the posterior brain.

**Figure 3.**
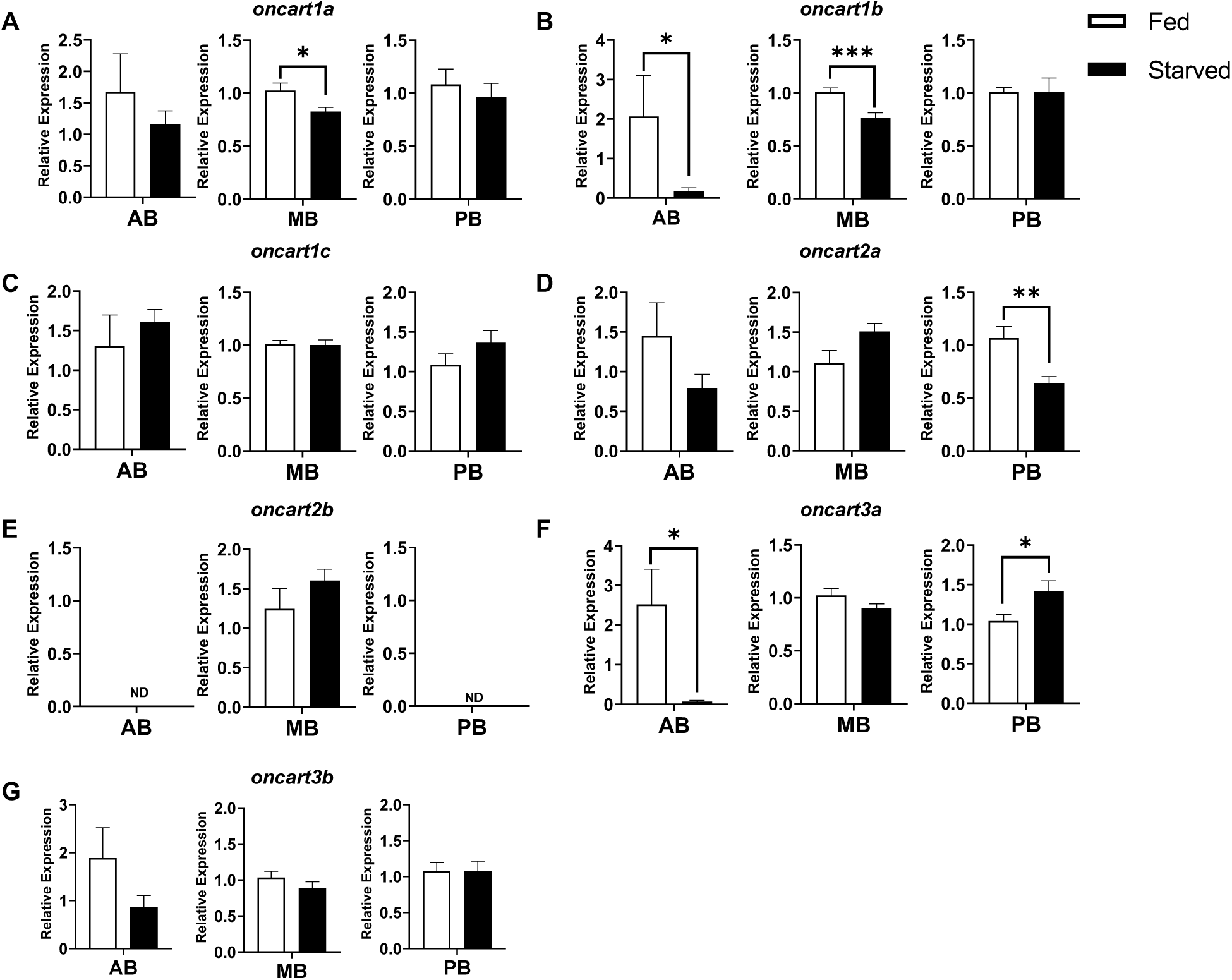
Relative expression of *cart* orthologues in major compartments of Nile tilapia brain following long term food deprivation (LD). Adult tilapia were not fed for 21 days. Expression of *oncart1a* (**A**), *oncart1b* (**B**), *oncart1c* (**C**), *oncart2a* (**D**), *oncart2b* (**E**), *oncart3a* (**F**), *oncart3b* (**G**) was analyzed in the AB, MB, and PB of the brain using qPCR (control n = 14, starved n = 14). Significant difference (p < 0.05) is indicated by (*), p <0.01 (**) and p<0.001 (***).

**Figure 4.**
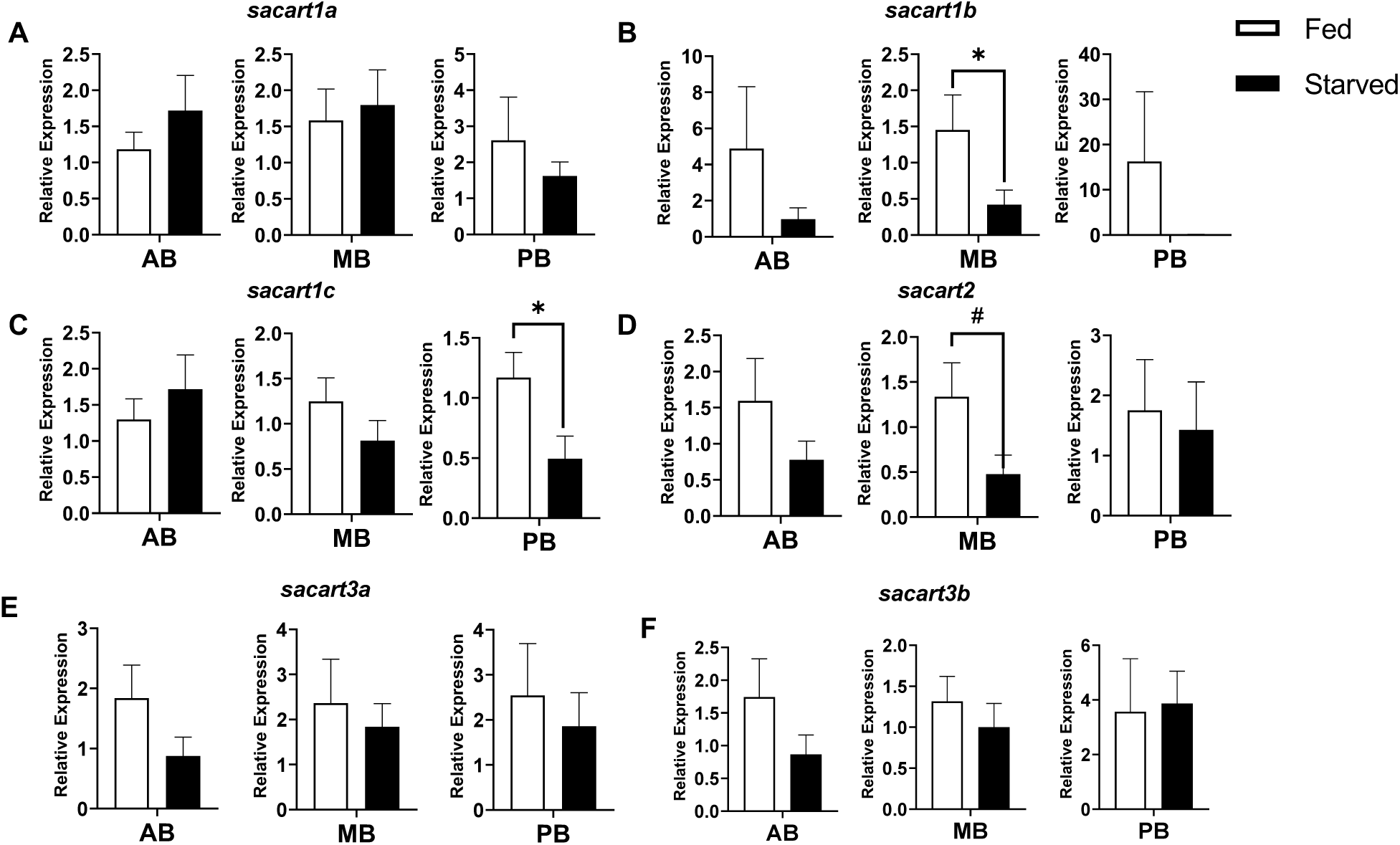
Relative expression of *cart* orthologues in major compartments of seabream brain following LD. Adult seabream were not fed for 7 days. Expression of *sacart1a* (**A**), *sacart1b* (**B**), *sacart1c* (**C**), *sacart2* (**D**), *sacart3a* (**E**), *sacart3b* (**F**) was analyzed in the AB MB, and PB of the brain using qPCR (control n = 9, starved n = 9). Significant difference (p < 0.05) is indicated by (*) and p = 0.070 (#).

## 4 Discussion

Fish have diverged from other vertebrate species in terms of long-term energy signaling (Hoskins and Volkoff, 2012; Meyer and Van de Peer, 2005) as in the case of leptin wherein its production is not affected by fat depots in fish (Blanco and Soengas, 2021). Moreover, the additional round of genome duplication in fish generated more copies of neuropeptides thus providing more complexity to central homeostatic regulation. Nile tilapia and gilthead seabream share homology in cart sequence, structure and distribution of expression, with the midbrain as the main site of their expression (Cabillon et al. 2025). However, in this work, we demonstrated that there are differences between these species in *cart* gene responsiveness to food deprivation. Following SD, *cart* expression in the midbrain significantly reduced for tilapia *cart1a*, *cart3a* and a trend to reduced *cart1b* (p = 0.085), as well as for seabream *cart1b*, *cart1c* and *cart3b*. Although there are differences in experimental setup and tissue sampling in the existing literature, the current consensus is that SD results to downregulation of *cart* genes (Bharne et al., 2013; Butt et al., 2019; Cai et al., 2015; Polakof et al., 2012). **Fig. 5A** illustrates that in four fish species with fully sequenced genomes, SD caused a decrease in expression in any of the piscine carts (Akash et al., 2014; Bonacic et al., 2015). Specifically, *cart1a* is SD-responsive in tilapia and zebrafish, while the other *cart1* genes are responsive in seabream and Senegalese sole (*Solea senegalensis*; Bonacic et al., 2015). *cart2* transcripts in both tilapia and seabream did not respond to SD, which is consistent with what was observed in zebrafish *cart2*(Akash et al., 2014). For *cart3* genes, tilapia *cart3a*, seabream *cart3b* and zebrafish *cart4* (clusters with *cart3a*; Cabillon et al., 2025) exhibited decreased expression in response to SD. Expression of *cart3* genes of sole did not change in response to SD (Bonacic et al., 2015).

**Figure 5.**
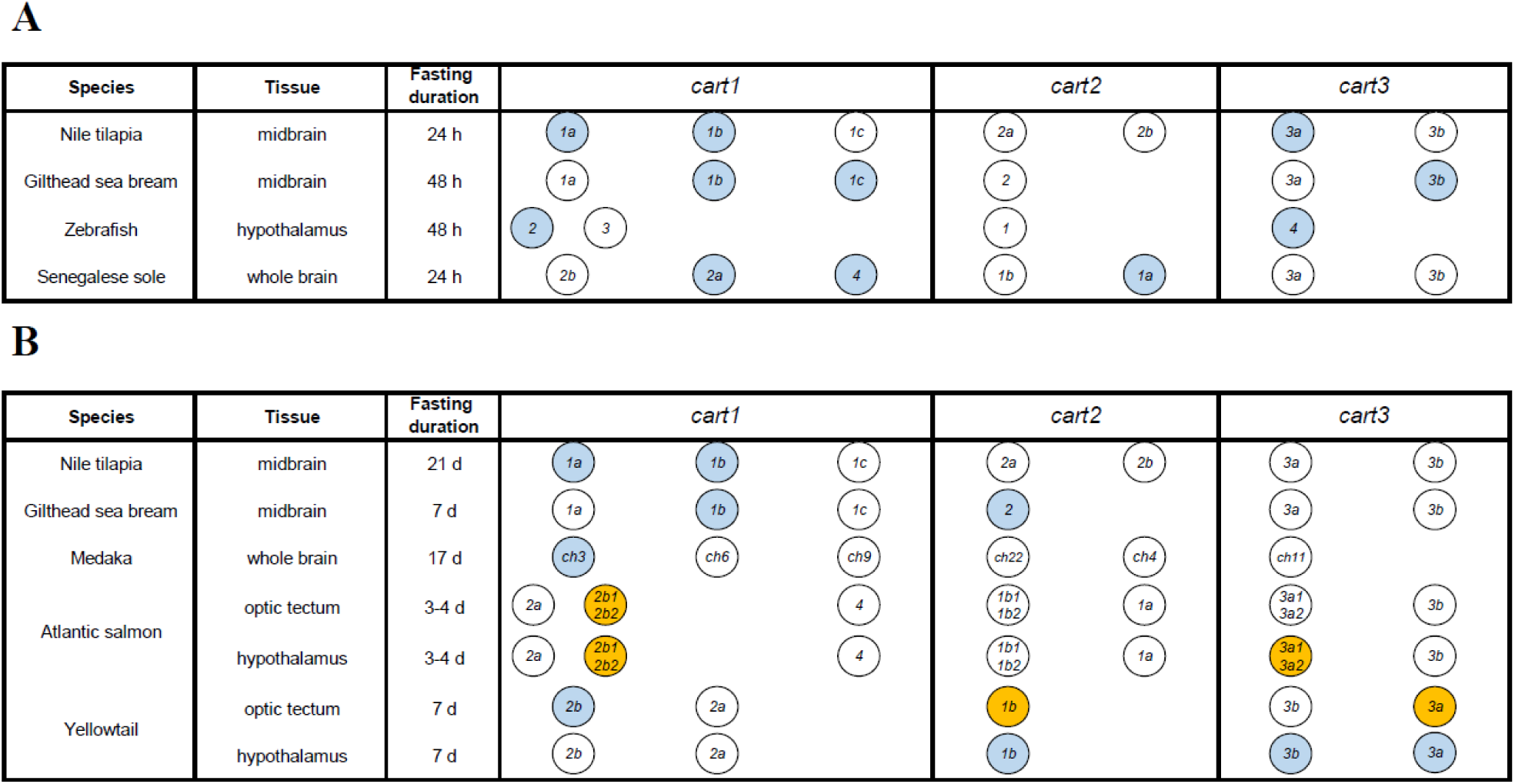
Responses of CART orthologs to food depravation in fish species with fully sequenced genomes. (A) CART genes of Nile tilapia, gilthead seabream, zebrafish and Senegalese sole in response to SD for either 24 or 48 h. (B) LD response of Nile tilapia, gilthead seabream, medaka, Atlantic salmon and yellowtail carts with varying durations. Blue-shaded circles indicate decrease in expression, orange-shaded circles indicate increase in expression and white-shaded circles indicate lack of significant response.

The effect of LD on *cart* genes expression is well-studied in mammals, birds and fish with a consensus that CART expression decreases after long-term fasting with some species showing no response to starvation. The responsiveness to food deprivation of CART is more varied in species with multiple orthologues such as chicken (1 out of 2) and medaka (1 out of 6) among others (Cai et al., 2015; Murashita and Kurokawa, 2011). As described for SD, the duration of feed restriction and the brain areas investigated in LD greatly varies according to the experimental system and animal. In addition, most fish species studied to date do not have fully sequenced genomes, and thus the complexity of the cart system in lower vertebrates might not be well-represented. In the current study, we identified a significantly reduced midbrain expression of specific *cart* genes in response to LD of Nile tilapia (*oncart1a*, *oncart1b*) and gilthead seabream (*sacart1b*, *sacart2*). This corroborate findings in other fish species. For example, LD led to decreased expression of *cart ch3* (herein clusters with *cart1a*) in the medaka brain and suppressed *cart2b* (herein clusters with *cart1a*) in the optic tectum of yellowtail (*Seriola quinqueradiata*; **Fig. 5B**). In the yellowtail hypothalamus, LD suppressed *cart1b* gene expression as in seabream *cart2* that clusters with it (Senzui and Fukada, 2023). Interestingly, yellowtail is the only species in which LD resulted in suppressed hypothalamic *cart3* (Senzui and Fukada, 2023). Evidently, *cart* genes that are involved in SD are not necessarily involved in LD suggesting some form of functional partitioning that can be influenced by genetic background, feeding strategy, metabolic state of the fish and other factors.

By analyzing *cart* expression in different brain regions, we showed that SD led to increased expression of *oncart1c* and decreased expression of *oncart1a* in the tilapia anterior brain. In seabream, SD resulted in decreased *sacart1c* in the same brain region. Similarly, SD in goldfish suppressed *cart* expression in the olfactory bulb (Volkoff and Peter, 2001). LD caused a decreased expression of tilapia *cart1b* and *cart3a* in the anterior brain which is in line with the results obtained in yellowtail wherein a decreased expression of *cart1b and cart2b* was observed in multiple regions of the anterior brain following seven days of starvation (Fukada et al., 2021). Contrastingly, the anterior brain expression of yellowtail *cart3a* and Atlantic salmon *cart2b* increased in response to LD challenges (Fukada et al., 2021; Kalananthan et al., 2021). Studies have shown that in the forebrain, neural input from the telencephalon and olfactory bulbs may affect feeding (Lin et al., 2000). More studies are needed to explain the differential expression of *cart* genes in the anterior brain in response to the energetic state of the animal and food availability. In the posterior brain, tilapia *cart1c* increased while seabream *cart1c* expression decreased following SD. LD supported reduced expression of tilapia *oncart2a* and seabream *sacart1c*. A similar decrease in the cerebellum was observed in yellowtail *cart2a* after LD (Senzui and Fukada, 2023). It was suggested that the satiety signals produced during feeding, primarily transmitted through the afferent fibers of vagus nerves from the upper gastrointestinal tract to the hindbrain in mammals may also be present in fish (Kalananthan et al., 2021; Schwartz et al., 2000). Further studies investigating different starvation durations and the response following re-feeding, are needed to determine whether functional partitioning of *cart* genes also exists across other species, and whether each gene exhibits a temporal threshold to energy depletion. Such insights would provide a more comprehensive understanding of the evolutionary divergence and specialization of each *cart* orthologue in relation to metabolic regulation and brain region.

The genes *oncart1a*, *oncart1b* and *sacart1b* exhibited anorexigenic functions as shown by the consistent decrease in expression in the midbrain following both SD and LD. We hypothesize that they have a major influence in the *cart*-specific appetite regulation in these species. We recently demonstrated that these cart peptides also possess similar proteolytic sites as mammalian Carts, which are not present in piscine cart2 or cart3 peptides (Cabillon et al. 2025). Consistent with previous works in various fish species, SD resulted in decreased expression of several *carts* in the midbrain for both studied species, yet the effect of SD on *cart* expression in other brain compartments was less consistent in Nile tilapia. LD, on the other hand, resulted in increased or decreased *cart* expression depending on the species and brain region. In conclusion, our data shows functional and temporal partitioning in a multigenic piscine CART system. Our findings further support that this partitioning is species-specific and therefore the identification of the key anorexigenic cart needs to be studied at the specific species of interest. Nevertheless, some functional conservation can be found as *carts* that were found phylogenetically and structurally close to mammalian Carts were starvation-suppressed and thus may exert a major influence in the anorexigenic function. Further studies using other fish species are needed to further understand how multiple carts regulate satiety in fish.

## 7 Author Contributions

Conceptualization – JB, AB, NARC, LK; Data curation – NARC, JB, AB; Investigation – NARC, LK, ASH, IMA; Methodology – AB, JB, NARC, LK; Formal analysis – NARC; Project administration – JB, AB; Supervision – JB, AB; Visualization – NARC; Writing (original draft) – NARC, JB; Writing (review and editing) – NARC, JB, LK, AB.

## 9 Acknowledgments

We thank Ms. Tatiana Slosman for her assistance in fish maintenance at Volcani Institute fish facility.

## 10 Data Availability Statement

The datasets generated for this study can be found in the GenBank according to the accession numbers detailed in the text.

